# Rapid and Highly Sensitive Detection of Fentanyl and its Analogs by a Novel Chemiluminescence Immunoassay

**DOI:** 10.1101/2023.11.15.567298

**Authors:** Jiang-yang Zhao, Mezbah Uddin, Daisy Unsihuay, William Butler, Terrance W. Xia, Jayson Z. Xu, Paul J. Jannetto, Ping Wang, Xiaofeng Xia

## Abstract

A highly sensitive immunoassay with sub-picogram limit of detection for fentanyl and a wide range of fentanyl analogs has been developed, using a novel high affinity antibody fused with NanoLuc, a small-size luciferase that can emit strong and stable luminescence. When applied to unaltered clinical human urine samples, the assay has sub-picogram limit of detection for fentanyl, with results fully concordant with gold standard liquid chromatography-mass spectrometry. When applied to environmental samples, the assay can detect as low as 0.25 pg fentanyl per inch^2^ environment surface. Assay turn-around-time is less than 1 hour, with inexpensive equipment needed and potential for high throughput automation or in-field screening. This novel assay has broad potential applications in clinical, environmental, occupational and forensic scenarios by detecting trace amount of fentanyl and its analogs to keep frontline workers safe.

## Introduction

Fentanyl and its analogs have become one of the leading causes of overdose deaths in the United States. A report published in 2023 by National Vital Statistics System shows that every day, approximately 150 individuals die as a result of overdoses caused by synthetic opioids, primarily fentanyl[1]. Although the overdose epidemic was initially linked to the over-prescription of opioids, recent reports suggest that the majority of opioid-related overdoses are caused by illicit fentanyl and/or fentanyl analogs, either alone or in combination with other narcotics[2]. These drugs are usually difficult to identify by end users, as narcotics suppliers increasingly resort to adulteration with fentanyl analogs due to their lower costs and shortage of heroin[3]. However, since their lethal doses are much closer to effective doses than traditional opioids[4], it is very easy for drug users to overdose.

Fentanyl is 50 to 100 times more potent than morphine and 30 to 50 times more potent than heroin, so the rapid identification of its presence is key to prevent overdoses[5]. As even small amounts of fentanyl can become aerosolized and weaponized[6], identification of fentanyl in trace quantities is crucial to combating the current widespread epidemic. During the 2002 Dubrovka Theater crisis, hostages were reportedly exposed to gas containing carfentanil which resulted in the death of more than 120 hostages as well as the rebels[7]. It has also increasingly become a concern among the frontline workers that they may be exposed to highly potent opioids without knowledge[8].

For field-portable detection of fentanyl, “point-and-shoot” methods like Infrared or Raman spectrophotometry are easy to use but lack the sensitivity for trace amount detection and are used for examination for bulk seized drugs[9]. Traditional gas and liquid chromatography mass spectrometry (GC-MS and LC-MS) require expensive instrumentation, relatively large sample volume and extensive sample pretreatment, resulting in the reduction in the overall assay throughput [10-12]. On the other hand, homogenous immunoassays [13] or lateral flow assays (LFA)[14] can easily detect fentanyl and its analogs from a few drops of human samples, have medium to high throughput, and are relatively inexpensive [15]. However, there is still a gap in limit of detection (LOD) between immunoassays and the gold-standard mass spectrometry methods, which could detect sub-nanogram to picogram per mL levels of fentanyl analogs from urine samples[10]. The LODs of currently available commercial immunoassays for fentanyl and its analogs are usually around 1 to 2 ng/mL, which are designed for screening of patients with overdose symptoms[15]. Such sensitivity is not sufficient for trace amount fentanyl detection from environmental sources with small sample volumes, or for long-term monitoring of occupational exposure which often result in sub-pathological levels of fentanyl in urine samples[16, 17].

To tackle this problem, we have developed a novel immunoassay with high sensitivity for fentanyl and its analogs, which is enabled by two major enhancements. The first enhancement is a novel rabbit monoclonal antibody with superior affinity and favorable binding kinetics against fentanyl analogs. The rabbit is a classic antibody production species, and the recent progress of rabbit monoclonal antibody (rb-mAb) technologies has enabled the development of high-affinity and highly specific rb-mAbs against a wide spectrum of antigens including very small sized molecules with picomolar affinities, compared to mouse-derived antibodies with nanomolar to sub-nanomolar affinities[18, 19]. The second enhancement is based on highly sensitive luminescence signals generated by NanoLuc. NanoLuc is an engineered luciferase originated from stabilizing the catalytic subunit of a deep-sea shrimp (*Oplophorus gracilirostris*) luciferase developed by Promega [20]. The 19.1 kDa luciferase is a monomeric protein which is highly soluble and stable, making it ideal for in vitro assays. NanoLuc can be recombinantly fused with antibodies to obviate the need for secondary antibody detection, thus simplifying the assay steps [21, 22]. Notably, another advantage of NanoLuc is its novel imidazopyrazinone substrate, furimazine, which can produce strong and stable luminescence with a half-life longer than 2 hours [20].

In this study, we first generated a high-affinity rb-mAb antibody for fentanyl and its analogs using a target-specific single-B-cell-based approach and screening immunoassays, then fused this rb-mAb with NanoLuc to combine its high affinity with the latter’s sensitivity to generate a fast immunoassay that can detect fentanyl below 1 pg/mL. We also demonstrated the utility of this high-sensitivity screening assay in microliter amounts of both unaltered clinical urine samples and environmental samples. Given its robust sensitivity and minimal requirement for sample amount, this novel assay has wide potential use in forensic, environmental, and occupational applications.

## Methods

### Animal immunization

Rabbit immunization was performed by Cocalico Biologicals (Stevens, PA) following a standard procedure [23]. All animal maintenance, care and use procedures were reviewed and approved by the company’s Institutional Animal Care and Use Committee (IACUC). New Zealand White rabbits were immunized subcutaneously with 500 μg of KLH-conjugated fentanyl (AAT Bioquest, Pleasanton, CA) emulsified with complete Freund’s adjuvant (CFA) for the first dose and incomplete Freund’s adjuvant (IFA) for subsequent boosts. The animals received 2 booster injections at 3-week intervals. The animals were sacrificed for spleen harvest 14 days following the final boost.

### Single B cell antibody cloning

The rabbit spleen was cut to small pieces with scissors. The tissue was then gently pressed through a 40-μm cell strainer using a syringe plunger. The cells were then laid on Lympholyte - Mammal (Cedarlane, Burlington, NC) and centrifuged at 800 g for 30 minutes. The lymphocytes were collected at the interface and washed before stained by biotinylated BSA-fentanyl and then AF647-streptavidin (BioLegend, San Diego, CA).

Single antigen-binding B cells were sorted using an S3 cell sorter (Bio-Rad, Hercules, CA), and deposited into a PCR tube with 5 μL lysis buffer (Takara Bio, San Francisco, CA). cDNAs were then prepared by reverse transcriptase (Takara Bio, San Francisco, CA) and the antibody coding regions were amplified by PCR. The PCR products were then cloned into an antibody expression vector with a CMV promoter.

### Recombinant antibody expression

HEK293 cells were transfected with the antibody expression vectors using electroporation [24]. After 5 days the cell culture medium was collected. To express the NanoLuc-fused antibodies, a DNA sequence encoding NanoLuc [20] was appended to the C terminus of the antibody light chain [21]. Cell transfection was done using electroporation. In screening assays, after 5 days the cell culture medium was collected and used in the immunoassays without purification. Purified antibodies including clone H23 were obtained by protein A chromatography.

### Antibody screening immunoassays and chemiluminescence immunoassay

To carry out the ELISA screening for antibodies, transparent 96-well plates were coated with 1 μg/mL BSA-conjugated fentanyl (AAT Bioquest, Pleasanton, CA). Cell culture medium was diluted 100 times and added to the wells to incubate for 1 hour at room temperature. The wells were washed extensively and HRP-conjugated goat anti-rabbit secondary antibody (R&D Systems, Minneapolis, MN) was added to incubate for 1 hour. After washing, TMB substrate (Thermo Fisher, Waltham, MA) was added and the absorbance was read with a plate reader.

To carry out the luciferase chemiluminescence assay, white opaque 96-well plates were coated with 0.2 μg/mL BSA-conjugated fentanyl (AAT Bioquest, Pleasanton, CA). The NanoLuc-fused antibodies were first incubated with the test samples containing fentanyl or its analogs for 15 minutes, then added to the coated wells and incubated at room temperature for 30 minutes. The wells were washed extensively and NanoLuc substrate (Promega, Madison, WI) was added to generate the luminescent signals. The signals were read with a plate reader (Agilent, Santa Clara, CA).

### LC-MS/MS method for fentanyl quantitation

In accordance with Clinical and Laboratory Standards Institute (CLSI) guidelines, a previously validated dilute and shoot method was used to quantify fentanyl in urine. Briefly, 100 μL of centrifuged clinical urine, standards, or controls were diluted 1:11 with isotopically labeled (deuterated, d5) internal standard for fentanyl in clinical laboratory reagent grade water and analyzed by an in-house developed LC-MS/MS method. 10 μL of sample was injected and analyzed by LC-MS/MS using one quantifying and one qualifying ion for each compound (fentanyl and the internal standard) with a total run time ∼8 minutes but capable of multiplexing on a TLX4 (Thermo Fisher, Waltham, MA) 6500 mass spectrometer (ABSciex, Framingham, MA). Fentanyl response curve was linear from 0.2-100 ng/mL. Intra- and inter-assay coefficients of variation (CV’s) was ≤10%. Method comparison to a reference method demonstrated slope of 1.0 ± 0.1 and R^2^ >0.98.

### Environmental test

To assess the application of the current method in environmental test, a 2-inch x 2-inch area of surface was spilled with indicated amount of fentanyl samples spiked into PBS buffer and allowed to dry. To perform the test, a cotton swab pre-wetted with sample buffer containing 10 mM phosphate buffer with 1% Tween 20, 1% Triton X-100 and 1% BSA was used to wipe the surface thoroughly. The swab was then soaked and swirled in 250 μL sample buffer to dissolve the sample. 50 μL of the sample was then used in the chemiluminescence immunoassay and the result was compared to the blank control to quantitate the presence of fentanyl.

### Clinical sample test

Deidentified clinical urine samples were collected following the protocols approved by the Institutional Review Board. For each test 2 μL urine samples were diluted with 50 μL of diluent (PBS with 1% BSA) and used in the luminescence assay. All samples were tested in duplicate.

## Results

### Antibody discovery and screening

A total of 616 clones were recovered from single fentanyl-binding B cell cloning, and recombinantly expressed by cloning into rabbit IgG backbone vectors and transfecting HEK293 cells. Competitive colorimetric ELISA was used to screen for fentanyl-binding antibody clones. The antibody-containing cell culture medium was diluted 100 times and added to the BSA-fentanyl coated wells, with or without 1 ng/mL fentanyl. The free fentanyl competes with the immobilized BSA-conjugated fentanyl, leading to reduced antibody binding to the surface. After incubation the plate was washed and incubated with an HRP-conjugated goat anti-rabbit secondary antibody. The fentanyl binding affinities of the antibodies were quantified by the inhibition percentage calculated using the following equation:

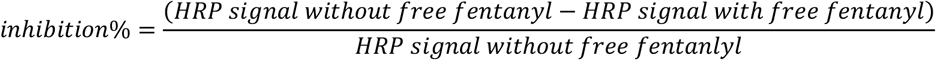

The binding of most of the antibodies to the immobilized BSA-fentanyl was markedly inhibited by 1 ng/mL free fentanyl (**Figure 1**), supporting their specific affinity for fentanyl. 23.9% (147 out of 616) of the clones showed inhibition rates greater than 90%, indicating high affinity binding.

**Figure 1.**
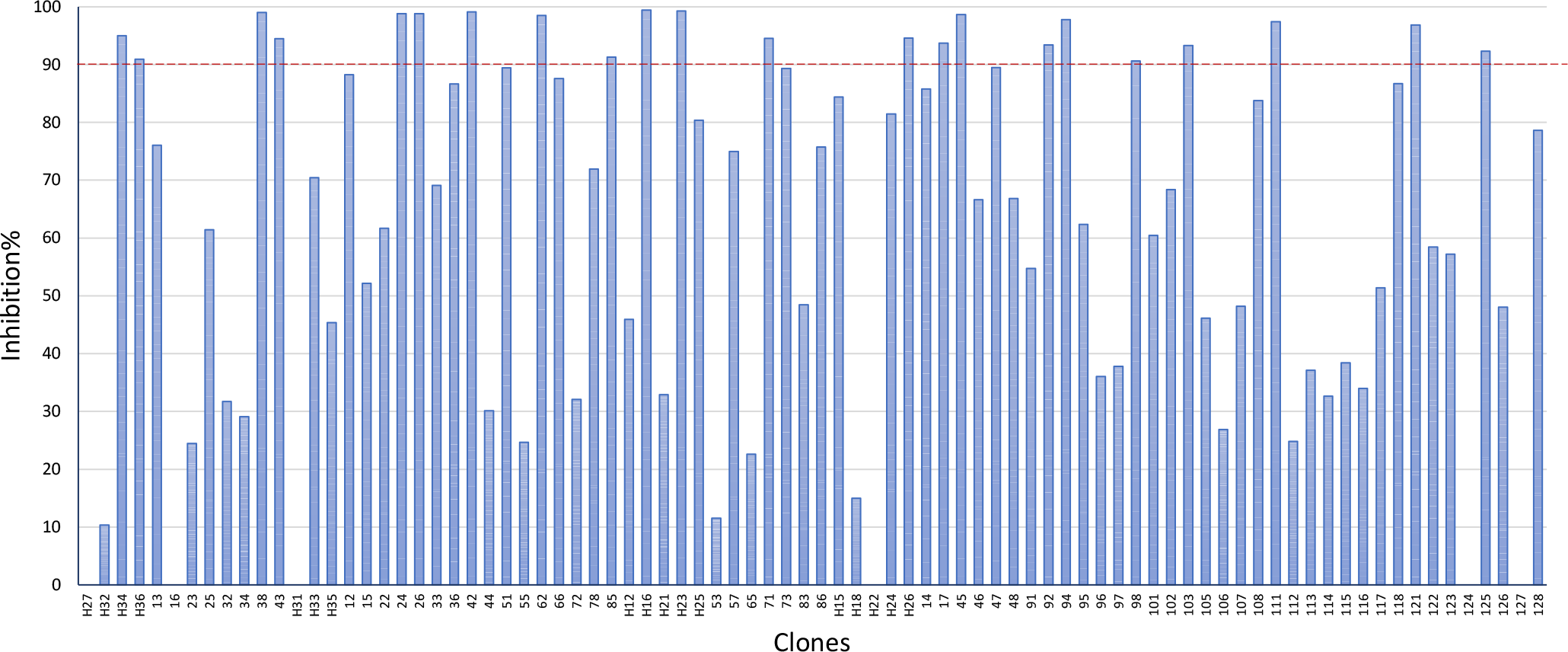
Screening of the clones. High affinity clones were selected that can achieve >90% inhibition with 1 ng/ml fentanyl. Only 84 of the 616 antibodies were shown for clarity.

### Antigen-antibody binding kinetics

To identify antibodies for rapid and sensitive detection of fentanyl, the time courses of the antibody-antigen binding were studied for 6 chosen antibodies (**Figure2**). The antibodies were added to BSA-fentanyl coated wells and incubated with indicated time. After that the plate was washed, the retained antibodies were quantified by an HRP conjugated goat anti-rabbit secondary antibody. The results showed that one of the clones, H23, exhibited fast binding kinetics and achieved more than 70% of maximum binding in 5 minutes. The rapid kinetics and the high affinity of this clone facilitated the development of rapid high-sensitivity fentanyl detection assays.

**Figure 2.**
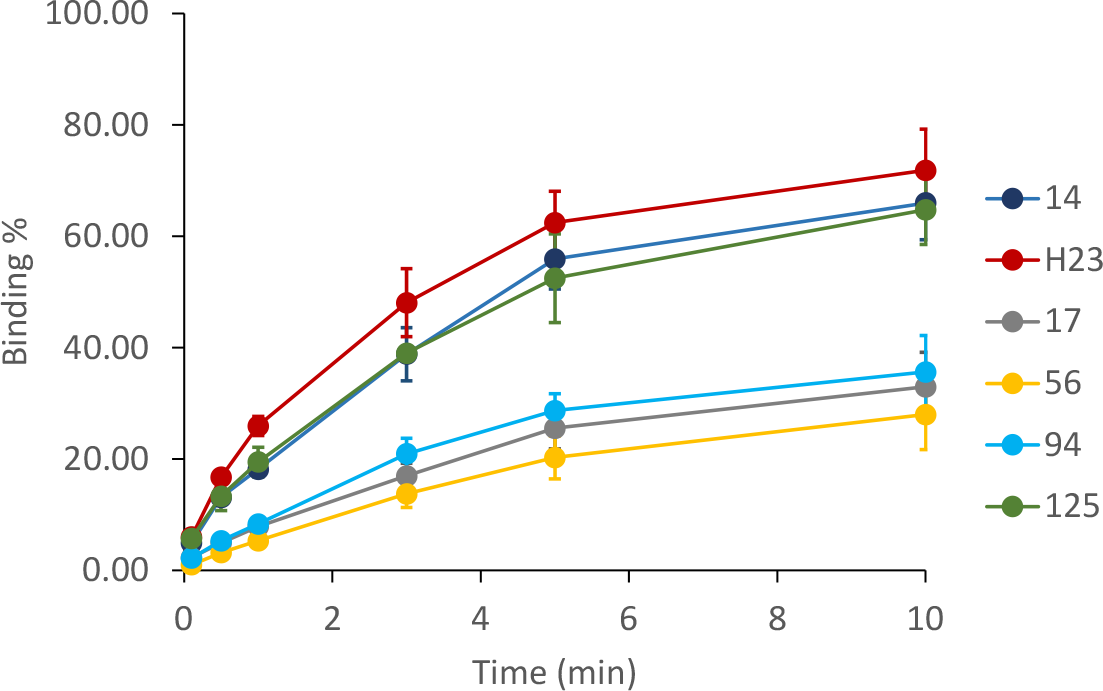
Antigen binding time courses of selected antibodies

### NanoLuc-fused recombinant antibody expression

To further simplify the assay procedure and increase the sensitivity, we tested whether the antibody could be expressed as a fusion protein with NanoLuc, so that it could directly generate intense signals without using secondary antibodies. The H23 clone was expressed as a NanoLuc fusion protein as described in Methods (**Figure 3A**). We achieved significant fusion protein expression, with the NanoLuc-fused antibody titer reached 15 μg/mL in the cell culture medium (**Figure 3B**). More importantly, intense luminescence signal was detected in the cell culture medium, indicating correct folding of NanoLuc when fused with the antibody. The result (**Figure 3C**) showed that even after 1 million-fold dilution, significant luminescence can still be detected in the medium, in contrast to the control medium expressed with antibodies without NanoLuc fusion.

**Figure 3.**
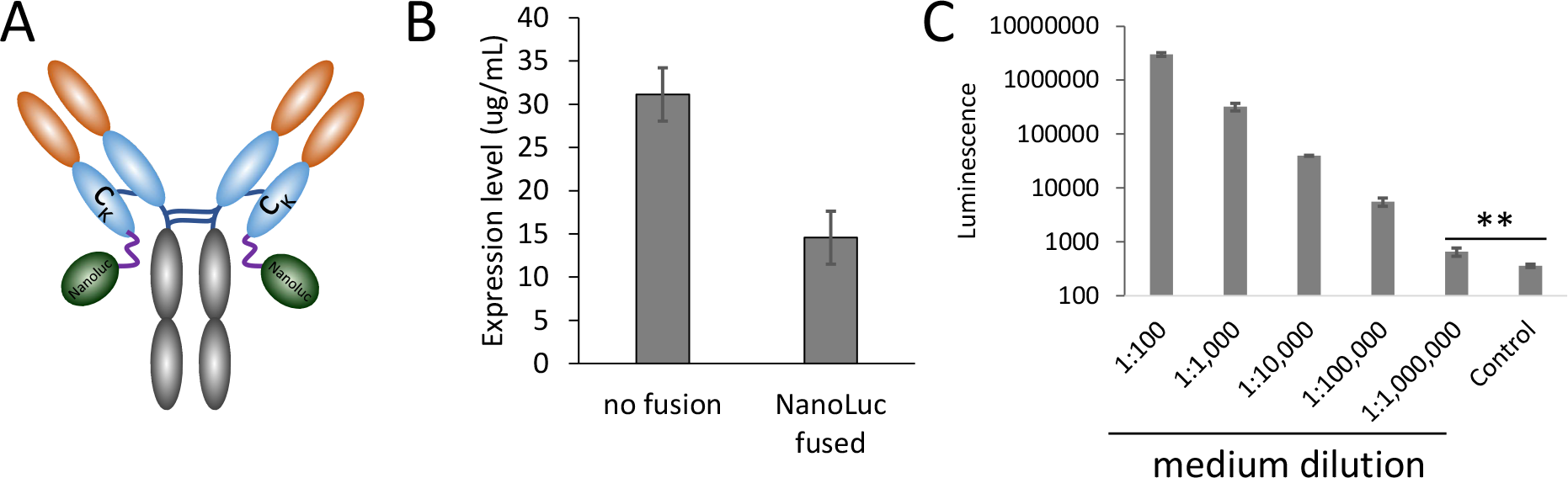
NanoLuc fused recombinant antibody expression

### Chemiluminescence immunoassay using the novel antibody

Using the NanoLuc-fused recombinant antibody H23, we developed a rapid chemiluminescence immunoassay method for fast and highly sensitive detection of fentanyl (**Figure 4A**), as described in Methods. Taking advantage of the fast binding kinetics of antibody H23, and the direct luminescent signal generated from the fused NanoLuc without using conjugated secondary antibodies, the assay can be completed within 1 hour. When the assay was carried out on samples spiked with fentanyl, the results (**Figure 4B**) showed that the assay can quantitatively detect fentanyl samples in both PBS buffer and human urine samples, with 50% inhibition (IC50) occurring at 15.5 pg/mL and 45.8 pg/mL, respectively. The limit of detection (LOD, defined as blank+3SD_blank_) of the assay in buffer and human urine samples was 0.25 pg/mL and 0.8 pg/mL in buffer and human urine, respectively, orders of magnitude lower than most reported GC-MS and LC-MS/MS (including UHPLC-MS/MS) assays, with LODs ranging from 5 to 500 pg/mL [12, 25].

**Figure 4.**
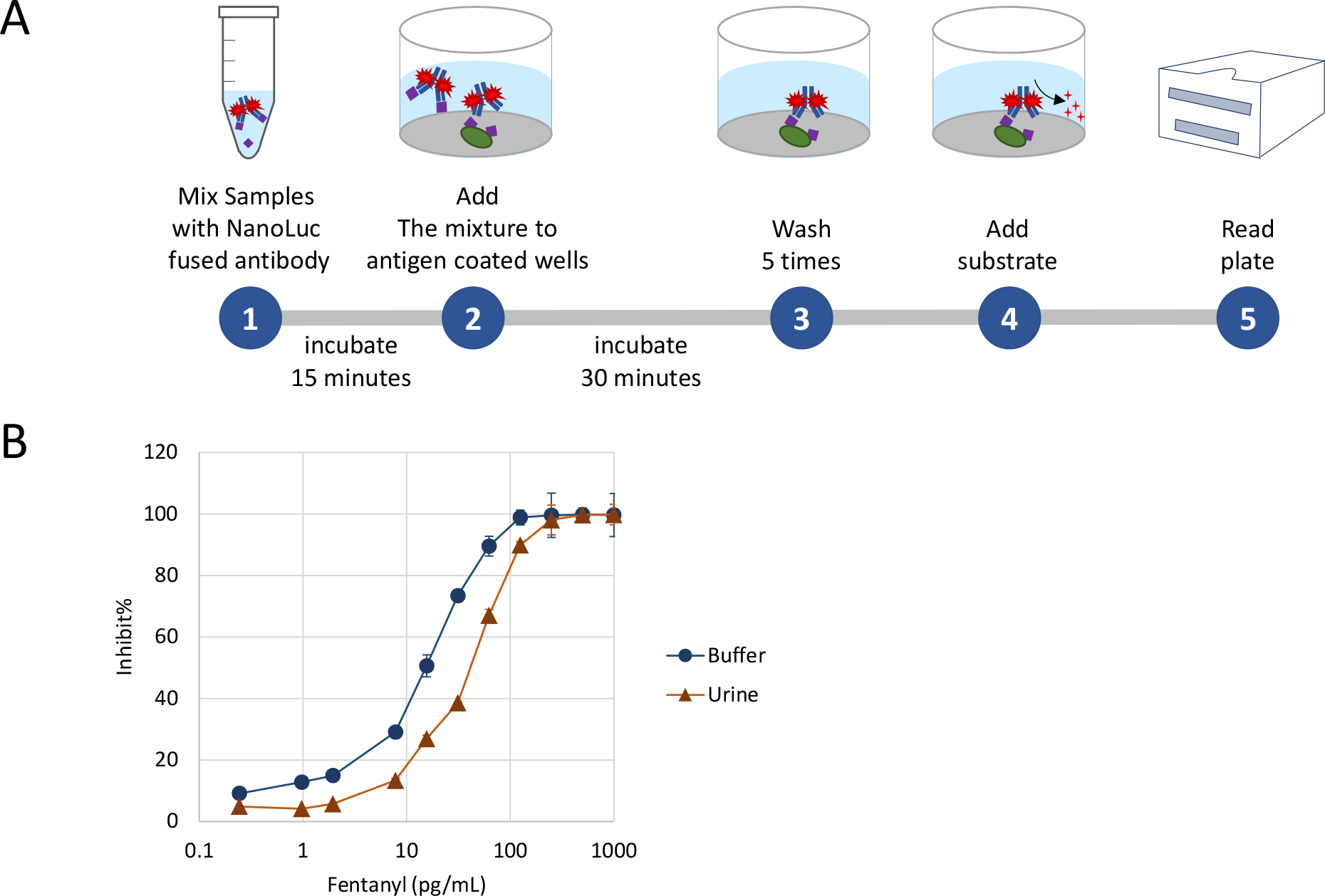
Chemiluminescence immunoassay for fentanyl detection

### Detection of fentanyl analogs

There are many existing fentanyl analogs and more keep emerging [26]. Therefore, it is important for screening assays to have broad cross reactivity for many analogs. A total of 17 different analogs were tested using our chemiluminescence assay, which was able to detect 14 of them (**Figure 5**). The assay sensitivities for the analogs (IC50 0.005 to 0.4 ng/mL) were comparable to that for fentanyl (IC50 0.05ng/mL). Notably, all the IC50 values are on par with the sensitivity of LC-MS/MS. However, the cost and time needed for our chemiluminescence immunoassay are much lower than that for LC-MS/MS. It is noteworthy that the assay was able to detect carfentanil with an IC50 of 0.4 ng/mL. Carfentanil is known as the most potent fentanyl analog, which is 10,000 times more potent than morphine.

**Figure 5.**
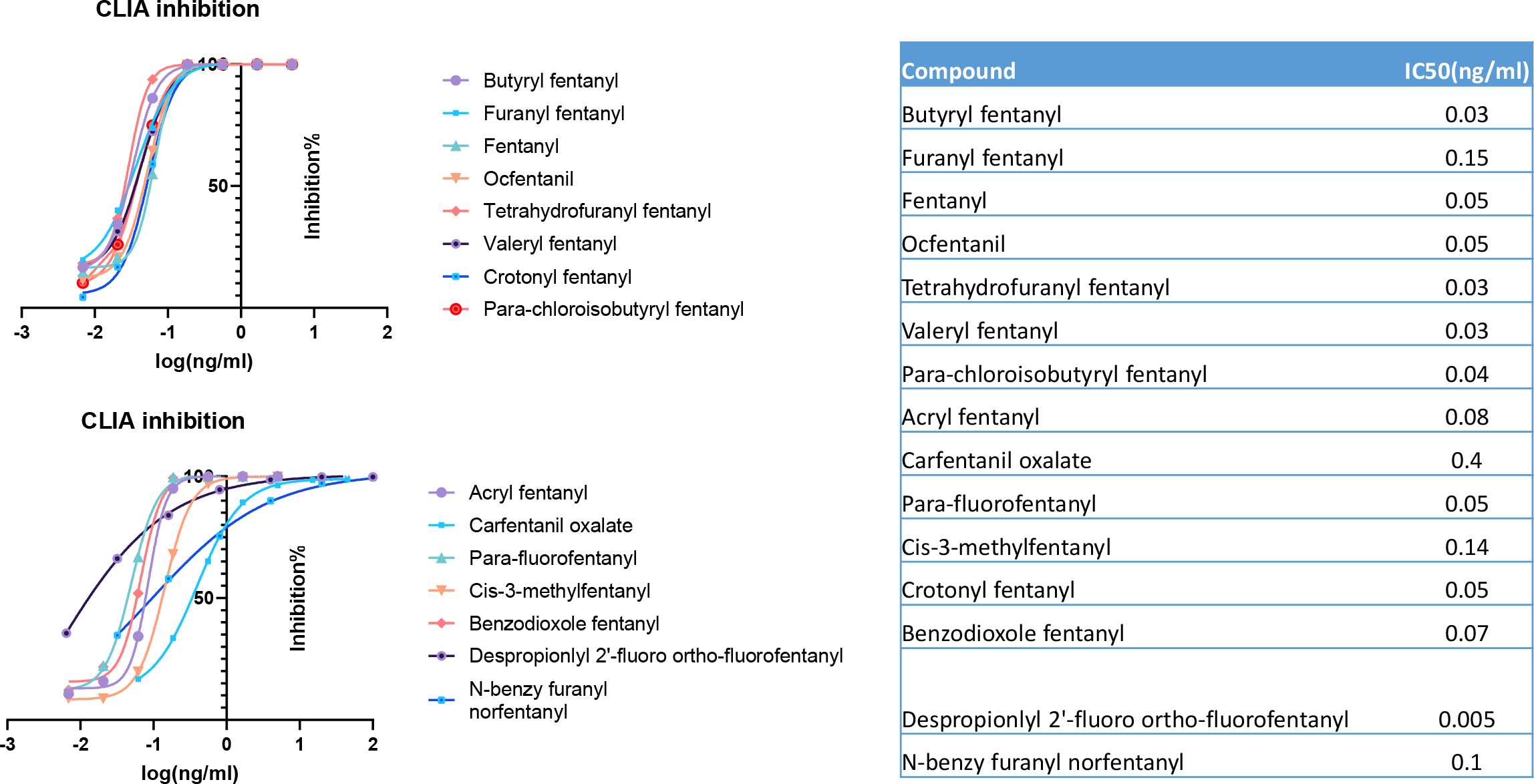
detecting fentanyl analogs

### Environmental testing using the new fentanyl chemiluminescence immunoassay

We then tested the utility of the new chemiluminescence immunoassay in detecting fentanyl from environmental samples. Trace amount of fentanyl was spilled on a fixed area and allowed to dry to mimic contaminated surfaces as mentioned in Methods. To collect the samples, the surface was thoroughly wiped with a cotton swab that was wetted with the sample buffer. The swab was then put into a tube containing 250 μL sample buffer to elute adsorbed chemicals by swirling the swab inside the tube (**Figure 6A**). Fifty μL of the sample was tested in the assay as illustrated in **Figure 4A**. The results (**Figure 6B**) showed that the assay successfully detected the lowest amount, 1pg fentanyl residue on a 2-inch x 2-inch surface, resulting in a lowest limit of detection at least 0.25 pg/inch^2^.

**Figure 6.**
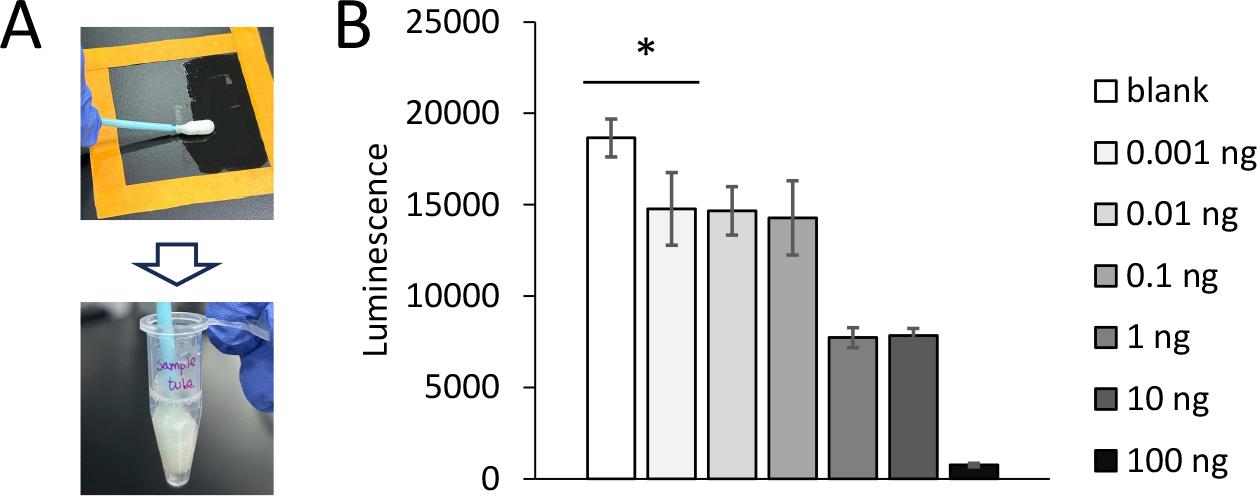
environmental test using the chemiluminescence immunoassay

### Clinical sample testing using the new fentanyl chemiluminescence assay

To assess the performance of the new fentanyl chemiluminescence immunoassay on clinical urine samples, especially those with low fentanyl concentrations as determined by current state-of-art, 10 clinical urine samples that contain 0.2 to 0.9 ng/mL fentanyl measured by LC-MS/MS were tested using the chemiluminescence immunoassay method. The results (**Figure 7**) showed that using only 2 μL urine samples diluted to 50 μL of final volume, the method was able to clearly detect the presence of picograms level of fentanyl in all urines. Notably, all the inhibition percentages were around 90% to 100%, which were near the upper limit of the linear range shown in Figs. 4 and 5, suggesting that the new chemiluminescence immunoassay had the potential to detect at least one to two-orders of magnitude lower concentrations of fentanyl from clinical urine samples, which would broadly expand the utility of this screening or presumptive assay.

**Figure 7.**
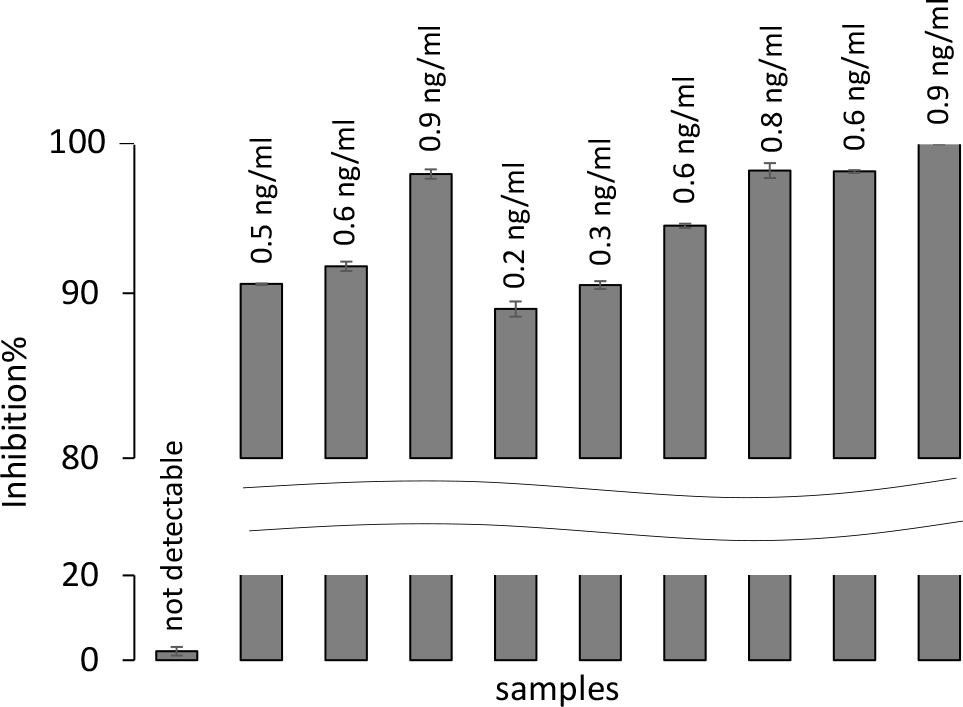
Detecting low concentration of fentanyl in human clinical samples, using the chemiluminescence immunoassay. The fentanyl concentrations determined by LC-MS are labeled above each sample.

## Discussion

In this study, we demonstrated the capability of a novel immunoassay to detect fentanyl and its analogs with very high sensitivity and very low sample volume requirement across multiple sample types, with potential for high throughput or in-the-field screening. Compared to existing immunoassays, this novel immunoassay can detect sub-picograms of fentanyl and analogs from environmental swabs and clinical urines.

In clinical or environmental field work, speed and ease-of-use of an assay are usually the most important among many considerations. Since our assay uses direct fusion of NanoLuc to antibody to generate luminescence, it avoids the secondary antibody binding and washing steps in conventional sandwich immunoassays. In addition, the high affinity and fast binding between the antibody clone H23 and fentanyl or its analogs allows short incubation time of 15 min at room temperature. Coupled with increasingly miniaturized portable plate readers[27], such assays can be deployed to the forefront of in-the-field screening. There is also great potential to leverage the novel antibody in other assay formats for improved ease of use. Lateral flow assay, or test strips, have been widely used for qualitative determination of many antigens, drugs, and narcotics which include fentanyl and its analogs, and has become popular among field workers and as a harm reduction strategy among drug users[9, 14, 28-30]. With the high-affinity antibody clones identified using our single B cell approach, coupled by novel materials to amplify the signal[28], it is possible to improve the current LOD of such assays by at least two orders of magnitude. Similarly, the improved antibodies may be used to develop highly-sensitive homogenous immunoassays to allow for high-throughput automation[15]. Another aspect of development is to make use of the strong stability and activity of NanoLuc, which may provide quantitative measurement when coupled with biosensor development and allow for quick quantification coupled with mobile devices. Currently, a linear response of NanoLuc signal spanning over three orders of magnitude with femtomolar LOD can be achieved by a prototype portable photomultiplier connected to a smart phone[27]. In the future, similar strategies of antibody discovery and assay development will broadly improve the detection sensitivity of many different narcotics.

Cross-reactivity to multiple fentanyl analogs is a desirable feature of screening and presumptive assays. However, false positives could be caused by known cross-reactivity to other non-opioids drugs like ziprasidone and risperidone, which share a core structure similar to fentanyl[31-33]. To solve this problem, one possible solution is to build a de-selection strategy into the antibody discovery process. Unlike traditional hybridoma technology or phage display, single B cell selection allows staining the antibody-expressing cells with multiple different targets simultaneously. Therefore, it is possible to assign a fluorophore to unwanted off-targets and eliminate the clones from the very beginning. Alternatively, one can also use similar animal immunization and antibody discovery strategy to identify ziprasidone and risperidone-specific antibodies and create a multiplex immunoassay panel. Such a panel will cover a range of structurally related chemicals and will be helpful to narrow down the plausible drug substance comprehensively early in the screening phase. Regardless of the method used, it is important to examine the structures of these chemicals and the strategy of hapten conjugation. It will be helpful to utilize different conjugation sites and linkers to expose the unique moiety to elicit desirable immune responses. Utilizing these conjugates, the antibody screening strategy used in our study can be easily applied to the discovery of many more drug-specific clones and the development of immunoassays with high sensitivity and specificity.

In the clinical diagnostic setting, the current commercially available screening immunoassays have a cut-off concentration at 1 to 2 ng/mL in urine[12, 15]. It should be noted that as low as 3 ng/mL fentanyl in blood may cause loss of protective airway reflexes and deaths[34, 35], and while most of the fentanyl is rapidly cleared from the body in an hour, only 10% is excreted in the urine[36, 37]. From this perspective, many drug users or front-line workers may have been exposed to pharmacologically effective or even dangerous levels of fentanyl with urine samples screened negative by the current screening immunoassays[38]. It is reasonable to postulate that overdose caused by more potent fentanyl analogs like carfentanil may lead to even higher rate of false negatives by these assays, and there is an emerging need to adopt more sensitive immunoassays for synthetic opioids[39]. Therefore, with sensitivity a thousand times higher than current screening immunoassays, the assay reported in this study has the potential to expand its use to clinical diagnostics in the future.

## Notes

### Competing Interest Statement

The authors have declared no competing interest.

